# The oncofetal protein IMP1 regulates the transcriptomic landscape to drive early events in pancreatic cancer progression and growth

**DOI:** 10.1101/2024.06.28.601273

**Authors:** Orçun Haçariz, Julia Messina-Pacheco, Elliot Goodfellow, Matthew Leibovitch, Andrew M. Lowy, Stephanie Perrino, Bertrand Jean-Claude, Zu-Hua Gao, Alex Gregorieff, Pnina Brodt

**Author notes:** Correspondence: Dr. Pnina Brodt, Research Institute, McGill University Health Centre, 1001 Décarie Blvd, Glen Site, Room E.02.6220 Montréal, QC H4A 3J1, Canada. UBC Department of Pathology & Laboratory Medicine, Vancouver, BC V6T 2B5. Authors share co-first authorship (Orçun Haçariz, Julia Messina-Pacheco).

## Abstract

**Background & Aims:** Pancreatic ductal adenocarcinoma (PDAC) has a dismal 5-year survival rate of 12% - the lowest of all malignancies. This is partially due to late diagnosis, as early stages of the disease, including the process of acinar to ductal metaplasia (ADM) are not presently detectable. Insulin-like growth factor 2 mRNA binding protein (IMP)1 is an oncofetal protein implicated in cancer progression. Here, we aimed to determine its role in the early stages of PDAC development and in the maintenance of the malignant phenotype.

**Methods:** IMP1 expression was analyzed in surgical PDAC specimens and in pancreatic tissue derived from KPC mice. Murine ductal organoids expressing the Kras^G12D^ mutant were treated with the IMP1 inhibitor BTYNB and RNAseq performed. The function of IMP1 targets was analyzed in an ADM model and the effect of IMP1 silencing on the growth of PDAC cells was evaluated *in vivo*.

**Results:** We found high expression of IMP1 in precancerous lesions of human and murine PDAC, but not in the normal pancreas. Blockade of IMP1 function impeded murine ADM and ductal organoid growth and profoundly altered the transcriptional landscape of the organoids, reducing the expression of cytokine-cytokine receptor interactors, cell adhesion and cell invasion mediators such as *Card11, Gkn3*, *Il13ra2*, *Mmp9*, and *Vcam1*. Gastrokine-3 and IL-13 in turn, enhanced the ADM process. Finally, IMP1 silencing in PDAC cells inhibited their metastatic outgrowth in mice.

**Conclusions:** IMP1 is a master regulator of early events in PDAC progression and a potential biomarker and target for this disease.

## INTRODUCTION

Pancreatic ductal adenocarcinoma (PDAC) is the most prevalent neoplastic disease of the pancreas, accounting for approximately 90% of all pancreatic cancers^1, 2^. The incidence of PDAC and PDAC-related death in the Western world are predicted to more than double in the next decade, rendering it the second leading cause of cancer-related death. With a 5-year survival rate in the USA of 12% and a median survival time after diagnosis of up to 12 months, PDAC is presently the deadliest of all common cancers^3, 4^.

One important aspect in understanding PDAC progression is the identification of precursor lesions, which are abnormal cellular changes that precede onset of malignancy. Pancreatic intraepithelial neoplasia (PanIN), intraductal papillary mucinous neoplasm, and mucinous cystic neoplasm, which differ in their histological characteristics, are the best-studied PDAC precursor lesions^5^. PanIN follows a histological stepwise progression that corresponds molecularly to a series of genetic alterations, including activating mutations in the oncogene *Kras* and the inactivation of the tumor suppressor genes *Cdk2na*, *Tp53*, and *Smad4/Dpc4* (reviewed in ^6^).

The exocrine pancreas consists of acinar cells that produce and secrete digestive enzymes, and ductal cells that line the pancreatic ducts. The acinar cells have a high degree of plasticity that is instrumental to maintaining tissue homeostasis and a capacity for regeneration in response to injury^7^. In response to triggers such as tissue injury, inflammation or stress, acinar cells can transdifferentiate and acquire a ductal-like phenotype, a process known as acinar-to-ductal metaplasia (ADM). The process entails a transcriptional downregulation of various genes including those encoding elastase, amylase, Mist1 and Cpa1, and the upregulation of ductal cell-specific transcripts including cytokeratin 19 (*Krt19*) and mucin 1 (*Muc1*) ^7–12^.

It has been well documented that PanINs can arise from pancreatic acini undergoing ADM ^13, 14^. The role of ADM in PDAC development was first demonstrated in mice by transgenic overexpression of transforming growth factor (TGF)-α ^15^ and can be replicated in 3D cell cultures when mouse acinar cell clusters are cultured in the presence of inflammatory cytokines or growth factors, or when they express activated Kras, leading to spontaneous transdifferentiation into duct-like structures ^9, 10, 15, 16^. Human acinar cells in 3D cultures can also undergo ADM in the presence of TGFβ ^17^. In patients with pancreatitis or other forms of pancreatic injury, ADM is transient and reversible ^18, 19^. However, in this metaplastic state, cells are susceptible to transformation by proto-oncogenes such as mutated *Kras*, which can drive formation of precancerous PanIN lesions^6^. With additional genetic hits in tumor suppressor genes or oncogenes, or exposure to environmental stress such as chronic inflammation, persistent ADM can progress from PanIN to invasive adenocarcinoma ^7, 20–23^. Suppression of trans-differentiation signals in cells undergoing metaplasia was shown to block progression to PanIN and PDAC^24^. Thus, identifying key players in the progression from ADM to PDAC can enhance our understanding of PDAC pathogenesis and lead to the development of novel, preventive and therapeutic strategies.

Insulin-like growth factor 2 mRNA binding protein (IMP)1 belongs to a family of three highly conserved oncofetal RNA-binding proteins that regulate the levels of target proteins post-transcriptionally by altering RNA localization, translation, and stability. During embryogenesis, the IMP paralogs (IMP1–3) play a role in stem cell renewal and organogenesis. After birth, their expression is absent or very low but they can be re-expressed upon malignant transformation ^25^. The main role of IMP1 in cancer cells is to impair miRNA*/*RNA-induced silencing complex (RISC)-directed mRNA decay by safeguarding the target mRNAs in cytoplasmic messenger ribonucleoproteins (mRNPs) ^26–28^. Several IMP1-regulated mRNAs encode proteins implicated in cancer progression including the transcription factors serum reponse factor (SRF) ^29^ and GLI; CD44, which plays a role in cell adhesion and invadopodia formation; integrin ITGB5; BCL2; c-Myc and IGF-2 (reviewed in^30^).

BTYNB (2-{[(5-bromo-2-thienyl)methylene]amino benzamide) (see **Supplementary Figure 1**) is a small molecule IMP1 inhibitor identified from a high throughput screen of a chemical library. It blocks IMP1 binding to target mRNAs, reducing their stability and protein levels ^31, 32^, and thereby reducing cell proliferation and colony formation of cancer cells ^31^. Ovarian cancer cells treated with BTYNB *ex-vivo* were growth-impaired *in vivo* ^29^.

High IMP1 expression levels in PDAC patients were associated with poor overall and progression-free survival^33^. However, the function of IMP1 in the initiation and early stages of the disease have not been elucidated. Here, we investigated the expression and role of IMP1 in PDAC progression using the IMP1 inhibitor (BTYNB), an animal model and clinical PDAC specimens.

## METHODS

### Human samples

With the approval of the institutional ethics review board of the McGill University Health Center, PDAC tissue samples were obtained from six patients with a history of chronic pancreatitis who underwent surgical resection at the McGill University Health Centre (MUHC). Formalin-fixed paraffin embedded (FFPE) tissue blocks that contain normal acini, ADM and PDAC areas on the same tissue section were selected for the construction of a tissue microarray (TMA). The clinical features of the six patients are shown in **Supplementary Table 1**. None of the patients received preoperative treatments, such as radiotherapy, chemotherapy, or biological treatments. Histology of all specimens were confirmed by two pathologists according to the WHO diagnostic criteria for PDAC ^34^.

### Mice

All mice experiments were conducted in strict accordance with the guidelines of the Canadian Council on Animal Care (CCAC) ‘‘Guide to the Care and Use of Experimental Animals’’ and under the conditions and procedures approved by the Animal Care Committee of McGill University (AUPs- MUHC-5260, 5733 and 7908). Unless otherwise specified, all mice used in this study were bred at the Research Institute of the MUHC (RI-MUHC) and used for the experiments at the ages of 7-12 weeks old. The origins of genetically engineered mice models (GEMM) and the induction of acute pancreatitis are described in detail in Supplementary Material. Necropsy information is provided in **Supplementary Table 2**.

### Synthesis of BTYNB

The IMP1 inhibitor BTYNB and the inactive control compound were synthesized by the Drug Discovery platform of the RI-MUHC, using the protocol descried in **Supplementary Figure 1**.

### In Vitro Models

#### Primary acinar cell isolation and three-dimensional culture

Primary pancreatic acinar cell isolation was performed as previously described ^35–37^, and detailed in Supplementary Material. The acinar cells were incubated at 37^0^C for 5 days with or without BTYNB (20 or 40 µM), a non-active control or vehicle (DMSO). Where indicated, recombinant murine IL-13 (Catalogue #210-13, ThermoFisher Scientific) and/or human Gastrokine 3 (GKN3, #CLPRO2033-2, Cedarlane, Canada) were added, with or without TGFα. Ductal structures were visualized on day 4 or 5 and bright field images of at least 10 fields were acquired for analysis as described in Supplementary Material.

#### Pancreatic ductal organoid culture

Primary pancreatic ductal cells were either isolated from the pancreas of LSL-*Kras^G12D^; CluCreERT; LSL-tdTomato* mice following Cre recombination induced by intraperitoneal injections of 200 µl tamoxifen (10 mg/ml) in corn oil for 3 consecutive days or induced *in vitro* by the addition of 0.5µM 4- Hydroxytamoxifen (4-OHT) in ethanol to the culture media. Pancreatic ducts were isolated as previously described ^38^ and detailed in Supplementary Material. Where applicable, the organoids were treated with BTYNB at the indicated dose for 48 hr, at which time brightfield images were recorded using the Lumascope720 microscope (Etaluma, Inc.).

### Cells

The pancreatic ductal adenocarcinoma LMP cell line originated from a liver metastasis that arose in the (*Kras^G12D/+^*;*LSL-Trp53^R172H^*^/+^;*Pdx-1^Cre^*) GEMM, as described in detail elsewhere^39^, and maintained in the Lowy laboratory since its derivation. In syngeneic B6.129 F1 mice implanted in the pancreas with LMP cells, tumor growth and metastasis mimic the aggressive clinical behavior of PDAC. The cells were routinely tested for common murine pathogens and mycoplasma contamination, as per the McGill University Animal Care Committee and the McGill University Biohazard Committee guidelines.

#### Generation and selection of IMP1-silenced LMP cells

Generation and selection of IMP-1 silenced LMP cells is described in detail in Supplementary Material.

### RNA sequencing

RNA sequencing was performed as described in detail in Supplementary Material. Briefly, pancreatic ductal organoids derived from ductal cells bearing the oncogenic Kras^G12D^ mutation were exposed to the IMP-1 inhibitor BTYNB, the non-active analogue or vehicle in triplicates. Total RNA of these cells were subjected to RNAseq. Validation for expressions of genes of interest by qPCR is described in Supplementary Material. Primer sequences are provided in **Supplementary Table 3.**

#### Meta-analysis

Downregulated DEGs (FDR < 0.05, log fold change < -1.5) were searched against the Cancer Genome Atlas (TCGA)/Genomic Data Commons Data Portal (https://portal.gdc.cancer.gov/) and the Pancreatic Cancer Database (http://pancreaticcancerdatabase.org/) ^40, 41^, using manual inspection for each gene.

### IMP1 binding to *Il13ra2* and *Gkn3* mRNA

Binding of IMP1 to *Il13ra2* and *Gkn3* mRNA was analyzed using RPISeq ^42^ and also biochemically by RNA co-immunoprecipitation (RIP) analysis, essentially as described elsewhere ^29^. The LMP cells were used as source of IMP1 and binding mRNAs. Immunoprecipitation was performed using a rabbit mAb to IMP1 (Cell Technologies,

#D33A2) and the Dynabeads Protein A immunoprecipitation kit (Invitrogen, #10006D), as per the manufacturer’s instructions. mRNA enrichment was assessed by qPCR.

### Statistical analysis

Two group comparison was performed by the Student’s t test or the Mann-Whitney test and multiple group comparisons (more than two groups) was performed by the One-way ANOVA, followed by Tukey’s multiple comparison test. The p values and the statistical tests used are specified, where applicable. Statistical analyses and graph generation were performed with GraphPad Prism (version 10.0.03).

## RESULTS

### IMP1 is highly expressed in human PDAC precursor lesions but not in the normal pancreas

To determine whether IMP1 may play a role in the initiating events and progression of this disease, we first analyzed a tissue microarray of six surgical PDAC specimens derived from patients with a history of chronic pancreatitis, where precursor lesions including ADM are seen, as well as FFPE sections derived from KPC mice. In surgical PDAC tissue, IMP1 was detectable in ADM lesions, PDAC and surrounding stroma, but not in the adjacent normal pancreas (**Figure 1A and B**). Staining intensity in PDAC was variable and ranged from weak or no staining to highly intense staining (**Figure 1C**). The increased IMP1 expression associated with ADM was confirmed by immunofluorescence performed on FFPE PDAC sections from one patient. In this specimen, IMP1 expression was confined to CK19-positive acinar cells in ADM and in α-smooth muscle actin (α-SMA) − expressing tumor associated stroma (**Figure 1D**).

**Figure 1:**
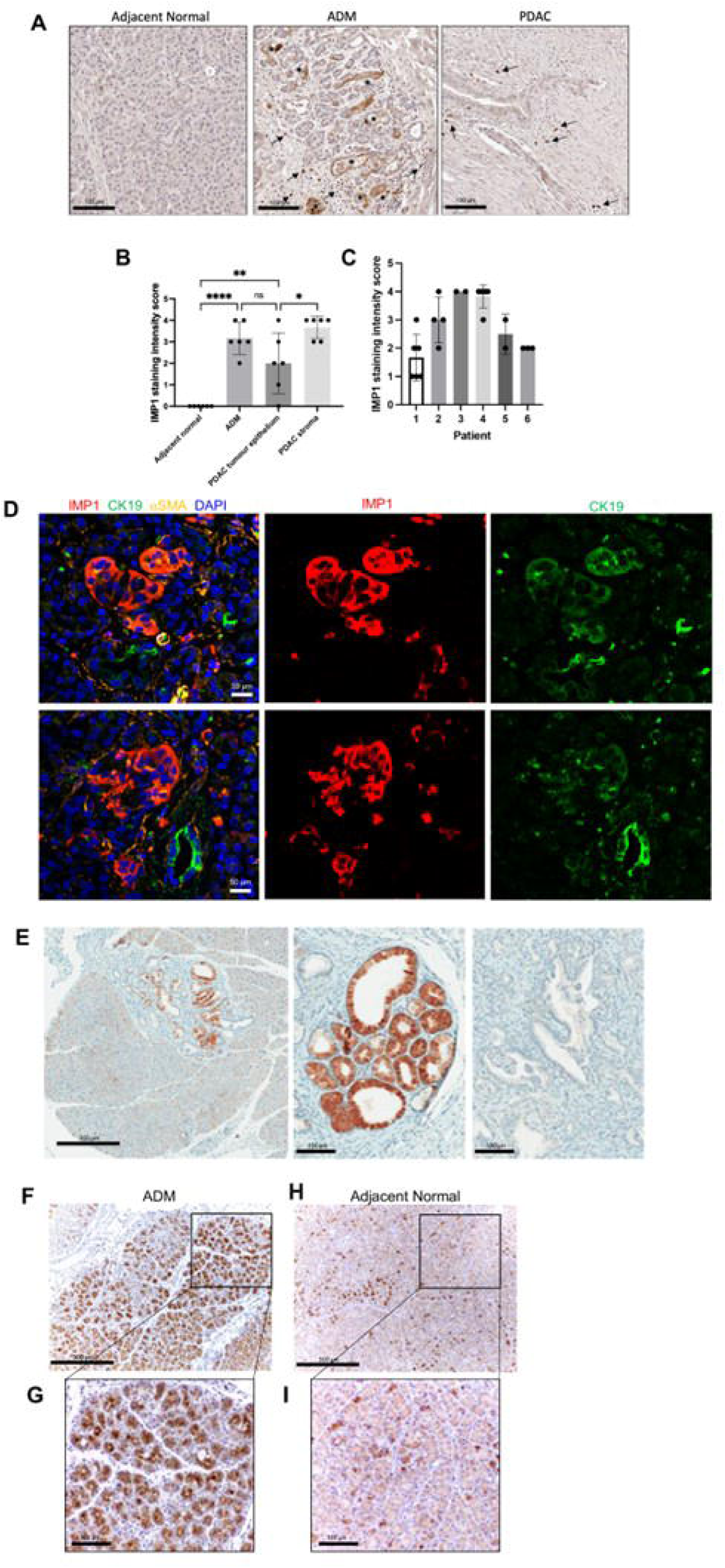
IMP1 is highly expressed in PDAC precursor lesions in both human specimens and mouse models. (**A**) Representative immunohistochemical images of IMP1 in a tissue microarray (TMA) of FFPE pancreas tissue from PDAC patients. IMP1 is highly expressed in ADM lesions (*) and the surrounding stroma of ADM and PDAC (arrows) but absent in adjacent normal parenchyma from the same patient. Scale bar: 100 µm. (**B**) Results of a semi-quantitative analysis of IMP1 staining intensity in different regions of the TMA showing a more intense staining in ADM and PDAC areas. n=6, one-way ANOVA with multiple comparisons. (**C**) Relative IMP1 staining intensity in PDAC tumors across patient samples. Each data point represents a core (1mm diameter) of PDAC tissue on the TMA. (**D**) Representative immunofluorescence images of human PDAC tissue, showing high expression of IMP1 in acinar cells that express the ductal marker cytokeratin 19 (CK19), a hallmark of ADM. Scale bar: 50 µm. *p < 0.05, **p < 0.01, ****p < 0.0001. Non-significant (n.s.) if p > 0.05. (**E**) Representative images of IHC of FFPE tissue obtained from KPC mice and immunostained with an antibody to IMP1. IMP1 is highly expressed in precursor lesions adjacent to PDAC, absent from adjacent normal parenchyma and variable in PDAC tumor epithelia. (**F-I**) Representative IHC stained pancreatic tissue obtained from a caerulein-treated mouse (enlarged in **G**) showing high IMP1 expression in ADM areas compared to adjacent normal areas of the same pancreas (enlarged in **I**).

### High IMP1 expression is also evident in precursor lesions of KPC mice and in caerulein-treated mice with acute pancreatitis

Similar results were obtained when we analyzed pancreatic tissue derived at necropsy from KPC mice (**Supplementary Table 2**). We found intense IMP1 staining in ADM and PanIN lesions, but not in the normal adjacent tissue. IMP1 was also detected at varying intensities in all cancerous tissue of KPC mice (**Figure 1E-I**).

Intraperitoneal injection of the cholecystokinin analog caerulein triggers acute pancreatitis in mice that mimics the pathology of the human disease. Chronic administration of caerulein, triggers ADM, immune cells infiltration and stellate cell activation, eventually leading to PanIN and PDAC when oncogenic activation occurs. We next evaluated the expression of IMP1 in ADM that developed in the setting of caerulein-induced inflammation. Immunohistochemistry revealed a marked increase in IMP1 expression in ADM lesions detected in FFPE sections of pancreatic tissue from these mice, but not from PBS (vehicle)-injected controls (**Figure 1G**), indicating that IMP1 was upregulated in the process of ADM regardless of the trigger.

### The IMP1 inhibitor BTYNB impedes ADM

We next sought to determine whether IMP1 was essential for the ADM process and ductal cell survival. Pancreatic acinar cells were cultured with 50 ng/ml TGFα for up to 5 days (**Figure 2A**) and confirmed that these cells underwent ADM (**Figure 2B and 2C**) that is associated with a decrease in amylase and a concomitant increase in cytokeratin 19 levels in the resulting ductal structures (**Figure 2D**). This process was inhibited when BTYNB, but not a non-active chemical analogue (control compound-CC) were added to the culture (**Figure 2B-D**).

**Figure 2:**
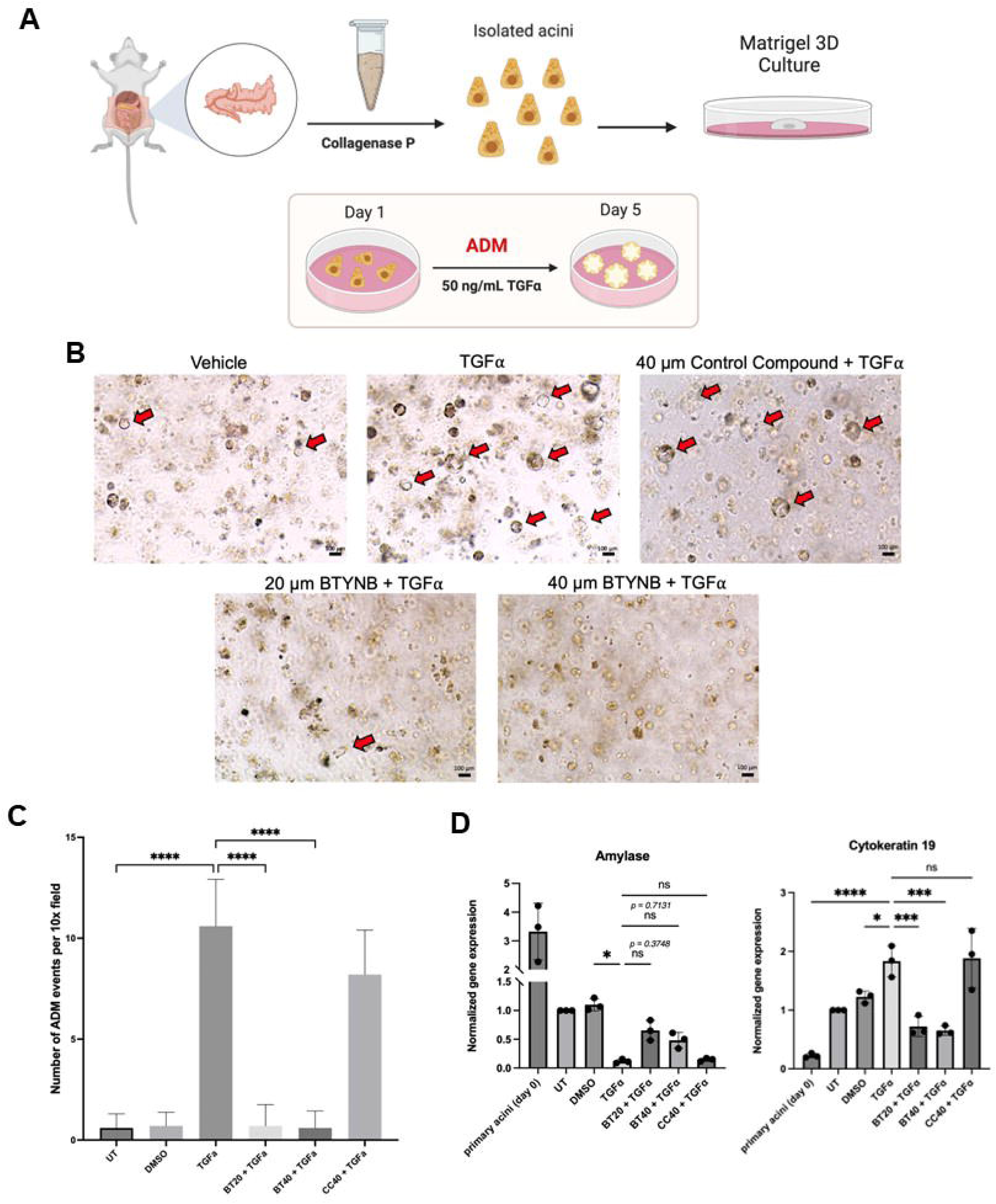
BTYNB, a small molecule inhibitor of IMP1, inhibits ADM in acinar cells treated with TGFα. (**A**) A schematic of the ADM assay. (**B**) Bright field images of ADM events showing a dose-dependent decrease in ductal structures (red arrows) in cultured mouse primary acinar cells treated for 5 days with TGFα (50 ng/ml) and BTYNB (20 µM or 40 µM), but not control compound. (**C**) Quantification of ADM events in the above conditions (n=3, 10 fields from each group, analyzed by one-way ANOVA with multiple comparisons). (**D**) Results of RT-qPCR. The addition of BTYNB reversed the increase in cytokeratin-19 expression to control levels but did not fully rescue amylase expression to baseline levels (n=3, one-way ANOVA with multiple comparisons). Values are presented as mean ± SD. *p < 0.05, **p < 0.01, ***p < 0.001, ****p < 0.0001, non-significant (n.s.) if p > 0.05.

### Inhibition of IMP1 by BTYNB also impedes proliferation and decreases survival of Kras^G12D^-expressing pancreatic ductal organoids

To determine if inhibition of IMP1 impacts ductal cell growth, we established ductal organoids derived from CluCreERT; LSL-Kras^G12D^;LSL-tdTom mice, where mutant Kras^G12D^ expression was induced by treatment of the cells with 0.5µM 4-OHTfor 48 hours. KrasG12D-expressing ductal organoids cultured in basal media with increasing concentrations of BTYNB displayed a significant reduction in size, as compared to untreated or control compound-treated organoids (**Figure 3A-C)**. Moreover, when EdU incorporation and cleaved caspase 3 levels were used to assess organoid cell proliferation and apoptosis in these organoids, respectively, a marked reduction in proliferation was observed in BTYNB-treated organoids relative to controls that was accompanied by increased apoptosis (**Figure 3C**).

**Figure 3:**
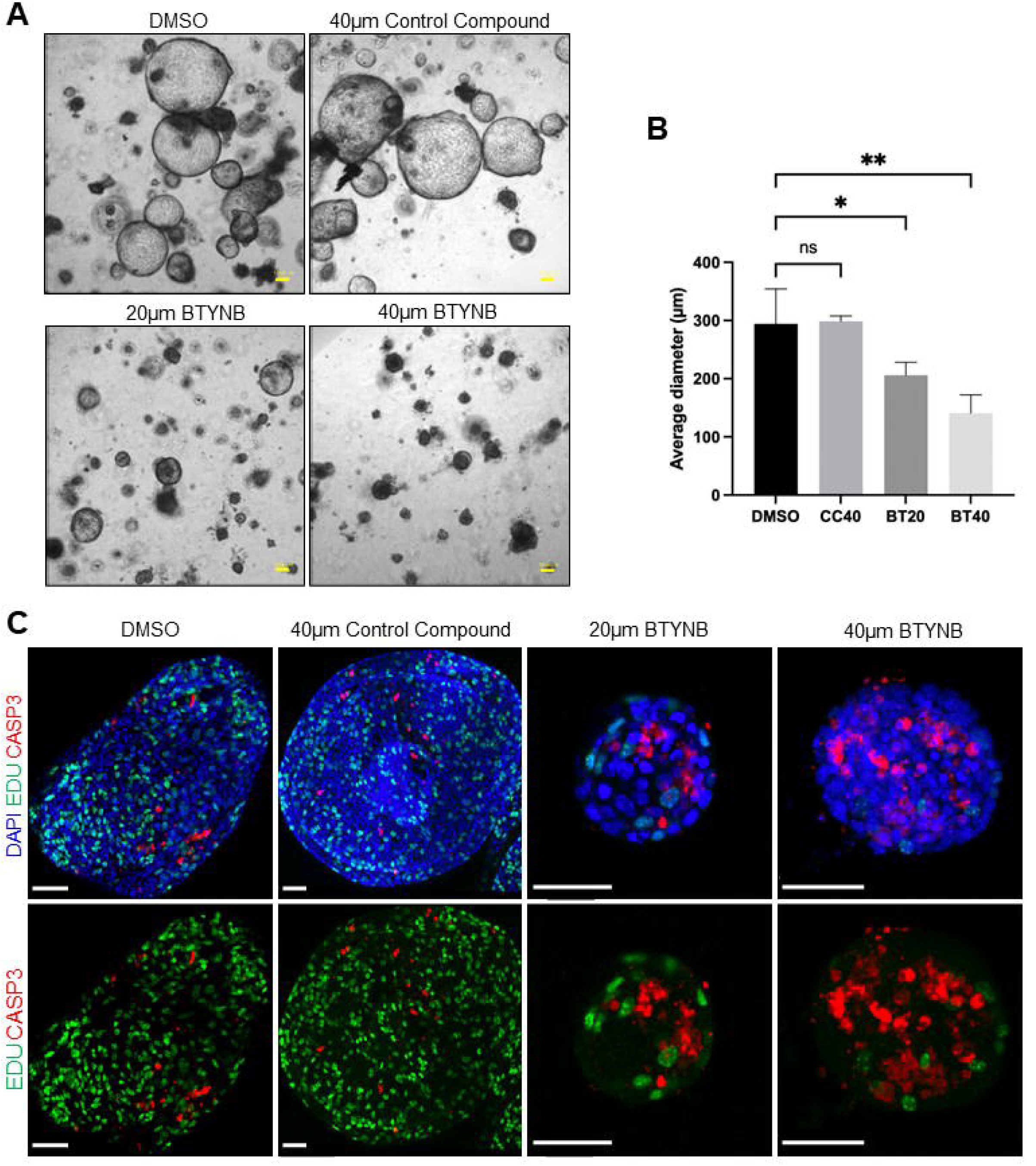
The IMP1 inhibitor BTYNB impairs proliferation and reduces survival of Kras^G12D^-expressing pancreatic ductal organoids. (**A**) Bright field images of Kras^G12D^-expressing pancreatic ductal organoids. A decreased size and impaired viability of organoids treated with (20 or 40 µM) BTYNB for 48 hours is evident. Scale bar: 50 µm. (**B**) Results of quantification of organoid diameters expressed as means ± SD (based on average of 10 4X fields per condition, n=3).*p < 0.05, **p < 0.01, Non-significant (n.s.) if p > 0.05 as determined using one-way ANOVA with multiple comparisons). (**C**) Immunofluorescence images of whole mount Kras^G12D^-expressing pancreatic ductal organoids following a 48-hour treatment with BTYNB (20 or 40 µM), vehicle (DMSO) or control compound. Scale bar: 50 µm.

### RNA sequencing identifies an IMP1-regulated transcriptomic profile in ductal organoids

To further identify mediators of early events in PDAC progression, we treated KrasG12D-expressing ductal organoids with 20 µM BTYNB (BT20), vehicle (DMSO) or the control compound at 20 or 40 µM (CC40, for increased specificity). Over 20 million pair-read sequences were obtained for each RNA sample. Principal component analysis showed a good separation between the control (CC40) and BT20- treated groups (**Figure 4A**). Further exploratory analyses showed almost identical results with the use of the vehicle (DMSO) (**Supplementary Figure 2**), indicating that CC40 was an appropriate control for BT20. We therefore used CC40 as control for RNAseq throughout this study. A total of 535 differentially expressed genes (DEGs) were identified when comparing BT20 and CC40-treated organoids (FDR < 0.05, absolute log fold change greater than 1) (**Supplementary Data 1**). Most of these genes (n=328) were downregulated in the BT20-treated group, as compared to the CC40- treated group (**Figure 4B**). Similarly, DEGs at high stringency cutoff (FDR < 0.00001, absolute log fold change greater than 3), including *Gkn3*, *Il13ra2*, *mmp9* and *vcam1*, were downregulated in BTYNB-treated organoids (**Figure 4C**). *Gkn3* was the most downregulated gene of all DEGs in respect to both log fold change and FDR value (log fold change: -9.3474, FDR: 7.12E-10).

**Figure 4:**
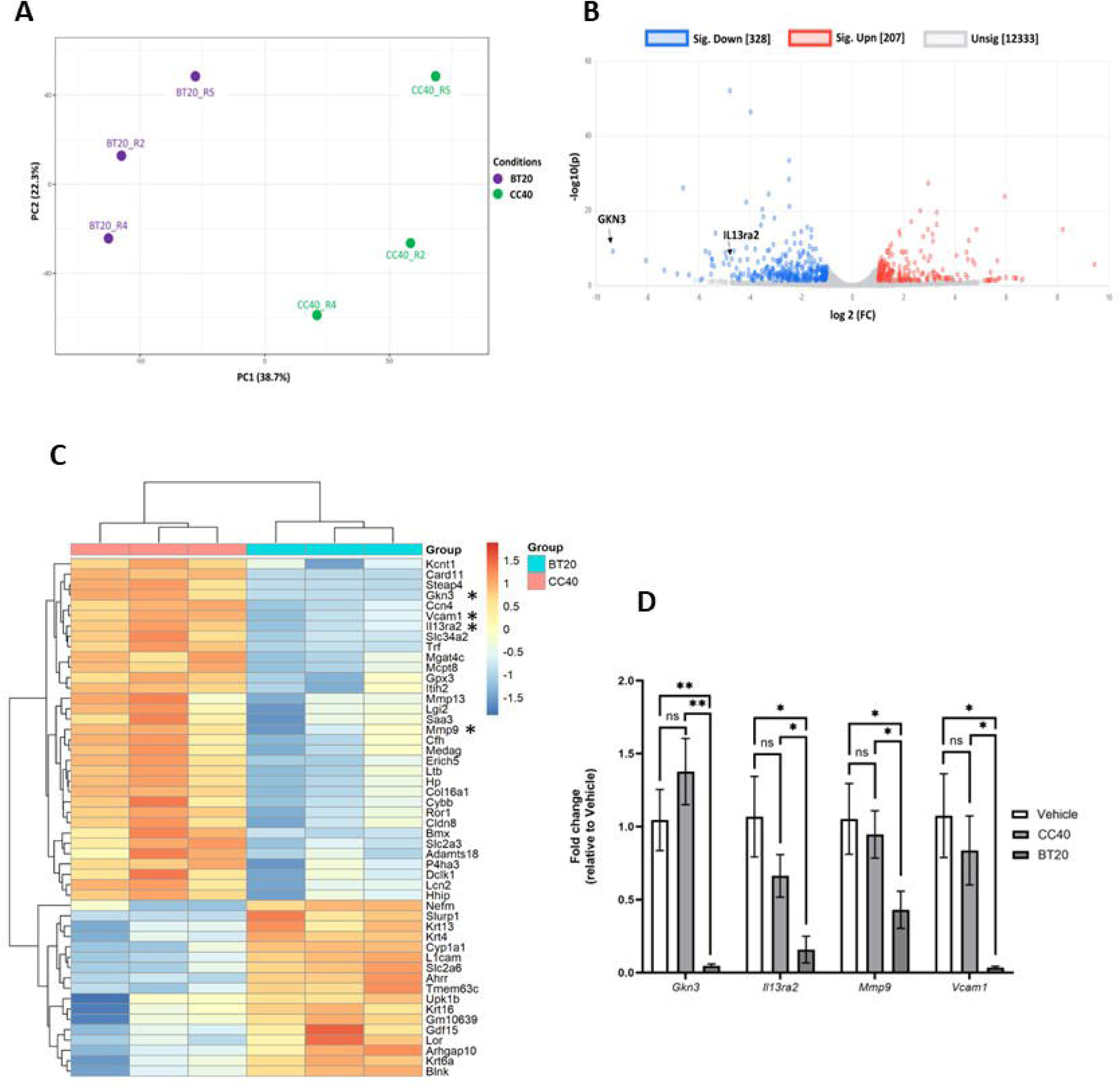
RNAseq analysis reveals an altered transcriptomic profile in ductal organoids treated with BTYNB. Kras^G12D^-expressing ductal organoids were treated for 48 hr with BTYNB or the control compound and RNA extracted for analysis (n=3). (**A**) Results of principal component analysis showing a clear separation between profiles obtained for BTYNB (BT20) and control compound (CC40)-treated organoids. (**B**) Volcano plots. The relative positions of *Gkn3* and *Il13ra2* transcripts are indicated, identifying *Gkn3* as the most downregulated transcript (log fold change = -9.3474, FDR < 0.00001). (**C**) A heatmap showing the top 50 DEGs in BTYNB, as compared to control treated organoids (FDR < 0.00001, absolute log fold change greater than 3), confirming that the majority of DEGs were downregulated in BTYNB-treated organoids. Genes of interest are indicated with an asterisk (*). (**D**) qPCR results confirming the downregulation of *Gkn3*, *Il13ra2*, *Mmp9* and *Vcam1* in BTYNB-treated organoids. Data are expressed as fold change relative to vehicle (DMSO). *p<0.05, **p<0.01 (Student’s t test, one-tailed).

Biological processes enrichment analysis of the DEGs identified various biological processes based on gene ontology (GO) annotation (**Supplementary Table 4**) that were related mainly to biologic regulation at the cell (e.g. cell migration and differentiation) and organ (e.g. organ morphogenesis and development) levels. Pathway enrichment analysis identified various physiology/pathology related pathways including chemical carcinogenesis, cytokine-cytokine receptor interactions and cell adhesion related pathways (complete data for the top 20 enriched pathways can be found in **Supplementary Table 5**). Heatmap analysis of the cytokine-cytokine receptor interactions pathway revealed downregulation of various cytokine and cytokine receptor genes (**Supplementary** Figure 3) and the most downregulated gene among them (log fold change -4.7098, FDR < 0.00001) was *Il13ra2*. The downregulation of several genes of interest was confirmed by qPCR, revealing marked reductions in the expression of *vcam1*, *gkn3 il13r*α*2* and *Mmp9* in BTYNB-treated organoids relative to controls (**Figure 4D**). Meta analysis revealed that among the genes downregulated in BTYNB-treated ductal organoids (FDR < 0.05, log fold change less than -1.5), caspase recruitment domain family member 11 (*Card11*), fibroblast growth factor receptors 2 and 4 (*Fgfr2* and *Fgfr4*), LIF receptor subunit alpha (*Lifr*), and several transcription factors (*Tbx3, Mycn*, *Maf* and *Creb3l1*), were identified by the Cancer Genome Atlas (TCGA)/Genomic Data Commons Data Portal within the cancer gene census category.

Of these, *Card11* was found with a high copy number alteration (39.43%, gain) in cancer cases in humans and it was markedly downregulated (>250 fold) in the BTYNB-treated organoids. Moreover, some of the downregulated genes including *Card11, Fgfr2, Fgfr4, Slc34a2, Il13ra2*, *Mmp9* and *Vcam1* were identified by the Pancreatic Cancer Database as upregulated (at the RNA and/or protein levels) in pancreatic cancer (**Supplementary Data 2**).

### IL-13 and Gastrokine 3 (GKN3) enhance ADM

Our analysis identified several IMP1- regulated transcripts not previously known to regulate the ADM process and ductal cell survival, including *Il13r*α*2 and Gkn3.* To determine the functional role of these mediators in acinar cell metaplasia, we used the ADM model and exogenously added IL-13 and/or GKN3 to acinar cells derived from normal mice, with or without TGFα. We found that the addition of IL-13 alone had little effect on the number of ADM events or ductal size. However, when added together with TGFα at a concentration of 50 ng/ml, IL-13 markedly increased the number of ADM and more significantly, the size of the individual ductal structures (**Figure 5A-C**). This increase in the number and size of ADM events was blocked with the addition of BTYNB (**Figure 5C&D**), implicating IMP1 in regulating downstream events in response to these mediators. Further confirmation of the important role of IL-13 in ductal cell outgrowth was obtained when ductal organoids were treated with 5 ng/ml IL-13. We observed a marked increase in the size of the organoids at 24 hr incubation with IL-13 and this was further amplified by 48 hr **(Figure 6A)**. Furthermore, qPCR analysis confirmed a significant increase in *Il13rα2* and *Gkn3* expression in organoids derived from Kras^G12D^ mice (**Figure 6B**), implicating mutant Kras in the regulation of these genes. We obtained different results when recombinant GKN3 (5ng/ml) was added to the ADM assay. Namely, similarly to TGFα, GKN3 added at 5 ng/ml significantly increased the number of ADM events **(Figure 6C)**, however, the ductal structures were smaller **(Figure 6B)** than those observed in response to TGFα or TGFα + IL-13 (**Figure 5C**).

**Figure 5:**
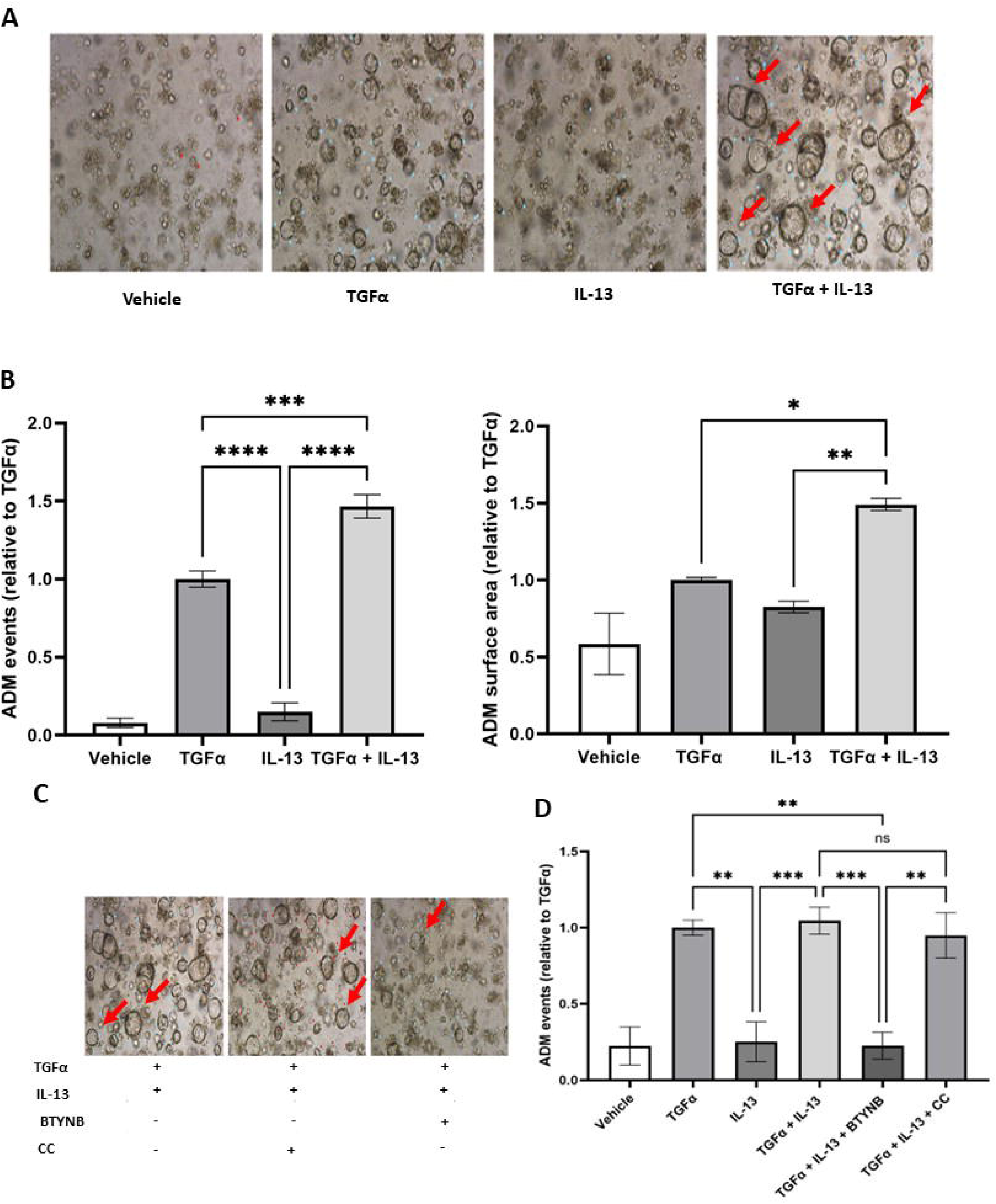
IL-13 increases the number and size of ADM events. The ADM assay was performed as depicted in Figure 2A. Acinar cells were incubated with 50 ng/ml TGFα with or without the addition of 5 ng/ml IL-13. (**A**) Representative images of ADM events, (**B**) results of quantification of ADM events following a 4-day incubation. (**C**) Representative images obtained in a separate experiment where the acinar cells were incubated with or without BTYNB or the control compound. (**D**) Results of the quantification of ADM events (5-day incubation, at which time some ADM structures in the IL13+TGF group collapsed due to excessive size). Results are expressed as a ratio to TGFα treated acinar cultures that were assigned a value of 1. *p < 0.05, **p < 0.01, ***p < 0.001, ****p < 0.0001, non-significant (n.s.) if p > 0.05 (As determined by one-way ANOVA; experiments performed twice each in duplicate cultures).

**Figure 6:**
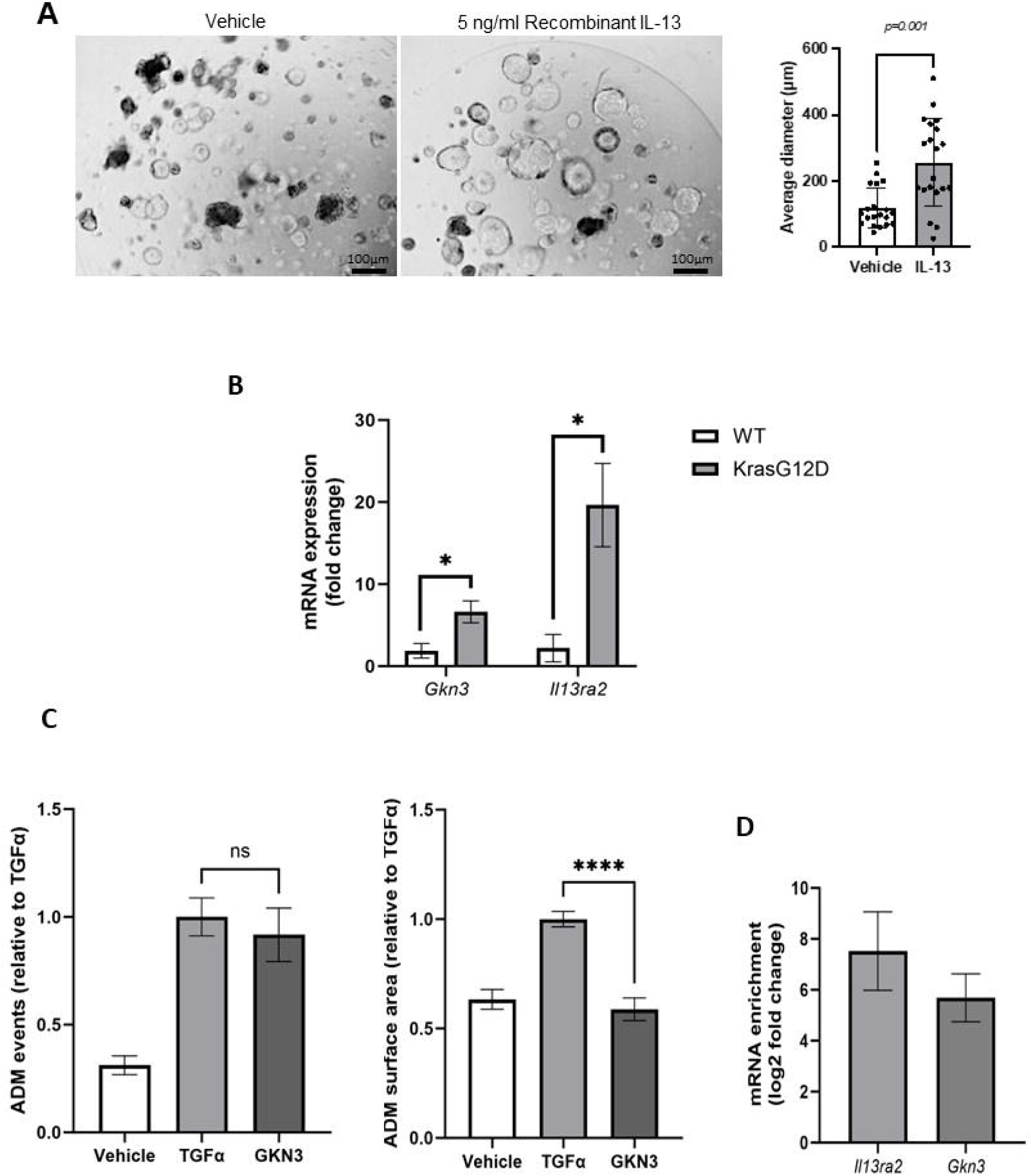
IL-13 and Gastrokine 3 promote the growth of ductal organoids and are direct targets of IMP1. Ductal organoids derived from wild type mice were incubated for 24 hr with 5 ng/ml recombinant IL-13. **(A)** Representative bright field images of the organoids treated (or not) with IL-13 for 24 hours and in the bar graph the mean organoid diameter based on 10 4X fields per condition and expressed as mean ± SD (n=3). **(B)** Results of qPCR analysis showing a significant increase in *Il13ra2* and *Gkn3* mRNA expression in KrasG12D ductal organoids as compared to wild-type organoids. *p < 0.05 (Student’s t test, two-tailed). **(C)** Results obtained after a 5-day culture of acinar cells with or without 5 ng/ml GKN3. The increase in the number of ductal structures was similar to the increase seen with 50 ng/ml TGFα. The results are expressed as a ratio to TGFα treated acinar cultures that were assigned a value of 1 (n=3). ****p<0.001 (As determined by one-way ANOVA). **(D)** Results of RNA co- immunoprecipitation performed on total LMP cell lysate using a Rabbit mAb to mouse IMP1 followed by qPCR performed on immunoprecipitated mRNA expressed as enrichment in *Il13ra2* and *Gkn3* mRNA transcripts in the eluate relative to the house- keeping gene HRPT1 that was used as a negative control.

The RPISeq tool ^42^ predicted a direct IMP1-*Il13ra2* mRNA interaction at probabilities of 0.75 (Random Forest classifier) and 0.9 (Support Vector Machine classifier) and of IMP1-*Gkn3* mRNA interaction at probabilities of 0.75 (Random Forest classifier) and 0.57 (Support Vector Machine classifier). To confirm empirically the direct interaction of IMP1 with these mRNA transcripts, we performed an RNA co-immunoprecipitation analysis using the murine PDAC LMP cell line and a rabbit monoclonal antibody to IMP1. A qPCR analysis revealed a significant enrichment in both *Il13ra2* and *Gkn3* mRNA in the immunoprecipitate of IMP1, as compared to mRNA of the housekeeping gene *HPRT1*, indicating that these transcripts were direct targets of IMP1 (**Figure 6D**). **IMP1 inhibition or silencing reduce the growth of PDAC cells *in vitro* and liver metastases *in vivo***. To assess the functional relevance of IMP1 to the growth of PDAC cells *in vitro* and their phenotype *in vivo,* we first treated LMP cells with increasing concentrations of BTYNB and analyzed their growth using the MTT assay. We found a significant reduction in cell proliferation at concentrations of ≥5µM BTYNB (**Figure 7A- B**), while the control compound had no effect, even at the highest concentration used (40 µm) (**Figure 7C**). In BTYNB-treated LMP cells, we confirmed a time dependent reduction in the levels of two known IMP1 targets namely, IGF-2 and c-Myc that are highly expressed in these cells, while, as expected, IMP1 levels *per se* were not affected (**Figure 7D**).

**Figure 7:**
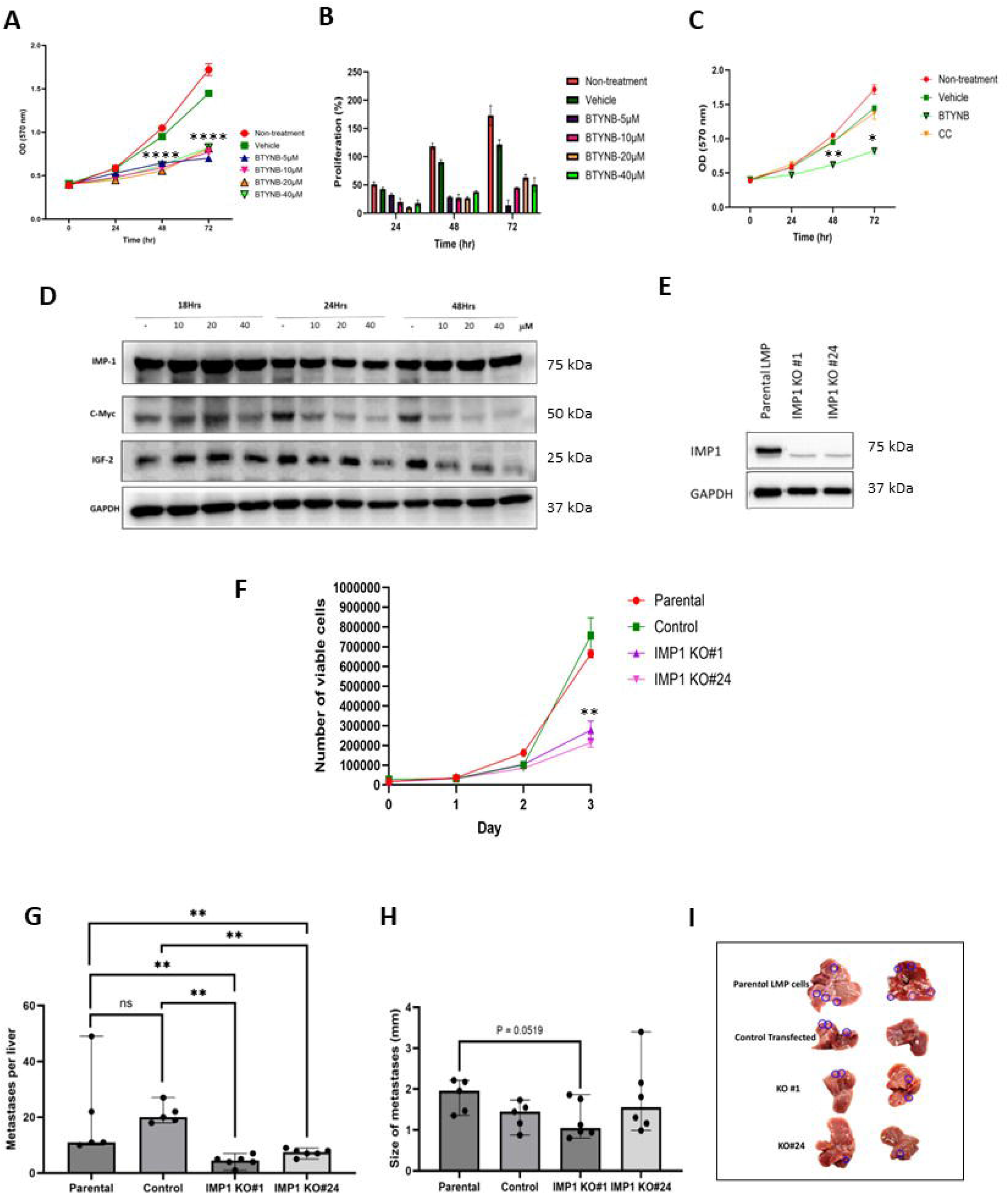
The IMP1 inhibitor BTYNB and IMP1 silencing inhibit the proliferation and liver metastasis of pancreatic ductal adenocarcinoma LMP cells. (**A-C**) Results of MTT assays expressed as OD measured at 570 nm (**A&C**) or as percent calculated relative to Day 0 (**B**) (n=3). *p<0.05, **p <0.01, ****p< 0.0001 (As determined by Student’s t test, two tailed). (**D**) Results of Western blotting performed on LMP cell lysate obtained after treatment with BTYNB for 72 hr showing reduced levels of the IMP1 targets c-Myc and IGF-2, as a function of BTYNB concentrations. (**E**) Western blotting confirmed reduced IMP1 levels in LMP cells following IMP1 silencing in LMP cells. (**F**) Results of cell proliferation assays analyzed using manual Trypan blue exclusion dye. (**G)** and **(H)** Medians (and range) of number and size of metastases per liver, respectively, analyzed 21 days following tumor cell inoculation via the intrasplenic/portal route. (**I**) Two representative livers from each tumor-injected group. Representative metastases are encircled. **p < 0.01 as determined by the Mann– Whitney test.

To test the role of IMP1 *in vivo*, we silenced IMP1 in LMP cells using a CRISPR-Cas strategy and compared the growth of two clonal populations with silenced IMP1 (**Figure 7E**) to that of wild-type and control transfected cells. We found that IMP1 silencing inhibited the growth of the tumor cells *in vitro* (**Figure 7F**) and significantly reduced the number of experimental liver metastases in syngeneic B6129F1 male mice injected with the cells via the intrasplenic/portal route (**Figures 7G and 7I**). A more minor reduction in the size of metastases (**Figure 7H**), suggested that a residual population within the IMP1 silenced LMP cells still expressed baseline IMP1 levels required for growth in the liver.

Collectively, our data identify IMP1 as a critical regulator of PDAC initiation, exerting its effect by controlling mediators of cellular metaplasia. They suggest, furthermore, that IMP1 is also involved in the maintenance of the malignant phenotype, identifying it as a potential therapeutic target in PDAC with high IMP1 expression levels.

## DISCUSSION

Our results implicate IMP1 as a mediator of early events in PDAC progression. We show here that IMP1 levels are elevated in human and mouse PDAC precursor lesions and can remain high in both the cancer and stromal compartments of PDAC tissue. We further demonstrate that blockade of IMP1 function by BTYNB impeded acinar to ductal metaplasia induced by TGFα *in vitro*, and also decreased proliferation and increased apoptosis of Kras^G12D^-expressing pancreatic ductal organoids. Moreover, treatment of aggressive PDAC cells with BTYNB reduced their proliferation, while IMP1 silencing in these cells markedly reduced their ability to form liver metastases. These results implicate IMP1 in multiple stages of PDAC progression, from early precursor lesions to acquisition of an aggressive malignant phenotype.

RNAseq analysis revealed a downregulation of multiple mRNA transcripts in IMP1- inhibited organoids. Among the transcripts most downregulated in BTYNB-treated ductal organoids were those of *Gkn3* and *Il3ra2*, as well as *Vcam-1* and *Mmp-9*. We have shown furthermore, that the addition of IL-13 to the ADM culture had a marked effect, increasing the number and size of the ADM events, in the presence of TGFα. This is consistent with several recent reports that identified IL-13 in ADM/PanIn lesions^43^ and IL13RA2/IL-13 as potential diagnostic and predictive biomarkers, as well as targets in PDAC ^44–47^. IL-13 / IL13RA2 signaling was also shown to trigger TGFβ1- induced fibrosis, suggesting that it could also play a role in the activation of a stromal response ^48^.

The role of GKN3 in metaplasia and pancreatic cancer initiation is less well understood. GKN3 is the latest of 3 stomach specific gastrokines to be discovered ^49^. It is expressed at relatively higher levels in cells during the gastric atrophy that occurs due to inflammatory conditions. Recent studies have strongly implicated GKN3 in gastric metaplasia. Thus, a recent single cell transcriptomic analysis identified *Gkn3* mRNA as a specific biomarker of mouse gastric corpus metaplasia ^50^. The role of GKN3 in human physiology and disease is less clear as the GKN3 protein could not consistently be identified in human gastric tissue, and an aberrant stop codon in the ancestral gene gave rise to the notion that GKN3 has evolved into a pseudogene ^51^. Menheniott et al suggested that the human GKN3 stop allele (W59X) could have been selected for among non-Africans due to its effects on pre-neoplastic outcomes in the stomach ^49^.

Of relevance is a recent study using lineage tracing and single-cell RNA sequencing that found that a number of canonical stomach spasmolytic polypeptide-expressing metaplasia (SPEM) markers including *Cd44* and *Gkn3* were enriched in a mucin/ductal subpopulation of ADM cells and that these cells also expressed *Gata5*, a transcription factor that is widely expressed in gastric/intestinal epithelium but not in the normal pancreatic epithelium ^52^. This identifies a hitherto unappreciated acinar-derived cell type in pancreatic metaplasia that shares a transcriptional program with gastric cells responding to injury and tumorigenesis ^52^.

In this context, our results identify *Gkn3* mRNA as a direct target of IMP1 and implicate this protein in promoting ADM. Thus, similarly to the findings in the gastric response to injury, inflammation and metaplasia ^50^, GKN3 may serve as a specific marker (and warning sign) and therapeutic target during early events in PDAC development.

We have shown here that *Gkn3* and *Il13ra2* mRNAs are direct binding targets of IMP1. The extensive transcriptional downregulation in BTYNB-treated ductal organoids suggest that some of the effects may be indirect and could be mediated via reduction in transcription factors or inflammatory mediators such as interferons ^53^ and TNF ^54^ that trigger chain reactions, leading to altered expression of other mediators. Indeed, MAPK and AKT signaling, as well as inflammation were identified as pathways most altered by inhibition of IMP1 activity in the ductal cells (**Supplementary Table 5**).

Among the genes downregulated in BTYNB-treated organoids were several extracellular matrix (ECM) proteins, in particular collagens XIIα1, XIVα1, and XVI1α1, tenascin and laminin, as well as metalloproteinases involved in ECM turnover such as MMP-9 and ADAM19. Interestingly, in mice with a targeted deletion in the IMP1 gene, downregulation of ECM proteins including collagen XIVα1 and tenascin were observed in the intestines and liver and were a major cause of impaired gut development and growth retardation ^55^. Our findings are consistent with these observations and position IMP1 as a major regulator of development and response to injury in the GI tract.

Of interest, several of the mRNA transcripts that were downregulated in BTYNB-treated organoids including *Card11, Fgfr2, Fgfr4, Slc34a2, Il13ra2*, *Mmp9* and *Vcam1* were identified as upregulated in pancreatic cancer by the Pancreatic Cancer Database (http://pancreaticcancerdatabase.org/) (**Supplementary Data 2**), suggesting that IMP1 may also orchestrate the transcriptomic landscape during the progression of the human disease.

Taken together, our data identify IMP1 as an important mediator of early events in PDAC progression, exerting its effect through regulation of multiple mRNA transcripts coding for proteins essential to metaplasia, ductal cell survival and transformation. IMP1 was identified in human colorectal cancer extracellular vesicles and we recently identified IMP1 in exosomes isolated from the plasma of PDAC patients (preliminary observation, data not shown). Moreover, IMP1 antibodies are already among a panel of antibodies to several tumor-associated antigens that are included in tailor-made panels for cancer detection ^56–58^. Given the critical importance of early detection for PDAC patients, our data and others identify IMP1 as a candidate biomarker that warrants further study, particularly in patients at high risk for the disease.

## Acknowledgments

This work was made possible by grants from the Cancer Research Society, the Canadian Institute of Health Research (CIHR- PJT-169032) and the Lustgarten Foundation to P. Brodt and grant # 390603 from the CIHR to A. Gregorieff. The authors of this manuscript are indebted to the Histopathology Platform of the RI-MUHC and its manager Ms. Fazila Chouiali for their help with immunohistochemistry, to Shibo Feng and Min Fu from the RI MUHC Molecular Imaging Platform for their technical expertise and help and to the McGill University and Génome Québec Innovation Centre for their excellent RNA sequencing service. O. Haçariz was the recipient of a RI-MUHC postdoctoral fellowship, J. Messina-Pacheco was the recipient of a Fonds de Recherche du Québec-Santé doctoral award (#305758) and E. Goodfellow was the recipient of a Cedars Cancer Foundation fellowship.

## Author contributions

PB was responsible for the conceptual framework of this study; PB, OH, JMP and AG contributed to experimental design and data analysis; OH, JMP, EG and ML contributed to data acquisition; PB, OH and JMP wrote the manuscript; EG was responsible for chemical synthesis; AG and AML contributed editorial comments; AML, BJC and ZHG provided resources.

## Data Transparency Statement

The authors declare that the data supporting the findings of this study are available within the paper and its supplementary information files. Unprocessed (raw) data can be made available by the corresponding author upon reasonable request. The RNAseq dataset (raw and processed) for this study is deposited in the NCBI Gene Expression Omnibus (GEO) database repository, with the accession number GSE248877.

## Abbreviations used in this paper

ADM: acinar to ductal metaplasia
BTYNB: 2-{[(5-bromo-2-thienyl)methylene]amino benzamide)
DEGs: differentially expressed genes
ECM: extracellular matrix
FBS: Fetal bovine serum
FFPE: Formalin-fixed paraffin embedded
GEMM: Genetically engineered mouse model
HBSS: Hanks’ Balanced Salt Solution
IMP1: mRNPs, messenger ribonucleoproteins
RISC: miRNA*/*RNA-induced silencing complex
RNAseq: RNA sequencing
IMP1: Insulin-like growth factor 2 mRNA binding protein
PanIN: Pancreatic intraepithelial neoplasia
PDAC: Pancreatic ductal adenocarcinoma
PBST: PBS-0.1% Tween
RI-MUHC: Research Institute of the MUHC
SRF: serum reponse factor
TGF: transforming growth factor
TMA: tissue microarray

## WHAT YOU NEED TO KNOW BACKGROUND AND CONTEXT

Pancreatic ductal adenocarcinoma (PDAC) is a deadly disease and there is an urgent need for improved early detection. The insulin-like growth factor 2 mRNA binding protein (IMP)1 is an oncofetal protein implicated in cancer development. We studied its role in PDAC initiation and progression.

## NEW FINDINGS

IMP1 is highly expressed in both human and murine PDAC precursor lesions and regulates the transcriptomic landscape of transformed pancreatic ductal cells. IMP1 targeting curtails metastatic outgrowth of PDAC cells.

## LIMITATIONS

The major IMP1-regulated transcripts in human and mouse PDAC cells or precursor lesions may be distinct and the human counterparts remain to be identified. The feasibility of detecting IMP1 in human serum requires confirmation.

## CLINICAL RESEARCH RELEVANCE

IMP1 upregulation was observed in surgical PDAC resections and IMP-1 targets identified by RNAseq are associated with PDAC progression, confirming its relevance to the human disease. IMP-1 and/or its regulated molecules are candidate biomarkers and potential therapeutic targets.

## BASIC RESEARCH RELEVANCE

This study integrates experimental and computational approaches to identify molecular mediators of early stages in PDAC progression. Our findings combined with the whole transcriptomic data provide insight into pathways regulating PDAC progression and a valuable resource to the research community.

