## Supplemental Material for "The oncofetal protein IMP1 regulates the transcriptomic landscape to drive early events in pancreatic cancer progression and growth"

### SUPPLEMENTARY MATERIAL

### SUPPLEMENTARY FIGURES

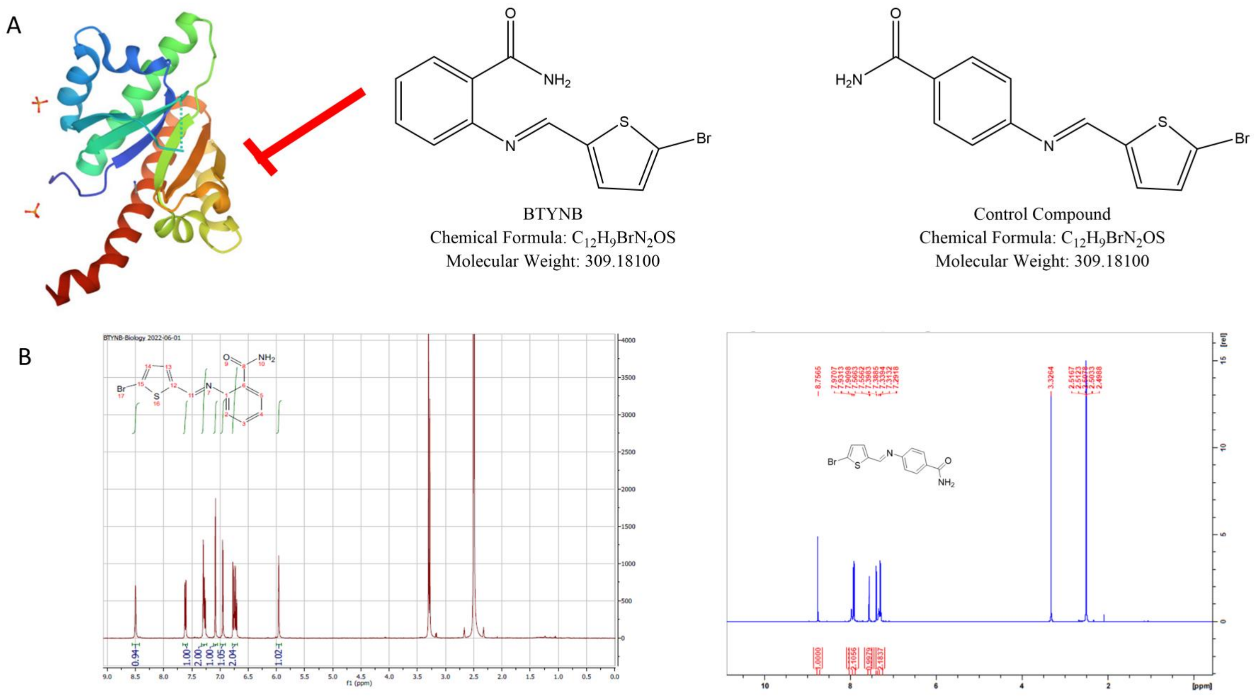


**Supplementary Figure 1. Synthesis and analysis of BTYNB and the control compound.** Shown in (**A**) are the chemical structures and in (**B**) the NMR profiles of BTYNB (left) and the control compound (right) synthesized by the Drug discovery platform of the RI-MUHC. HRMS of BTYNB, $C_{12}H_9BrN_2OS$, m/z: calculated $[M+Na]^+$ value is 330.9511 and observed $[M+Na]^+$ value was 330.9516: Δppm: 1.5 (<5ppm). MS of control compound: $C_{12}H_9BrN_2OS$ [M + H]: calculated value is: 308.83 and observed value was 310.83 m/z.

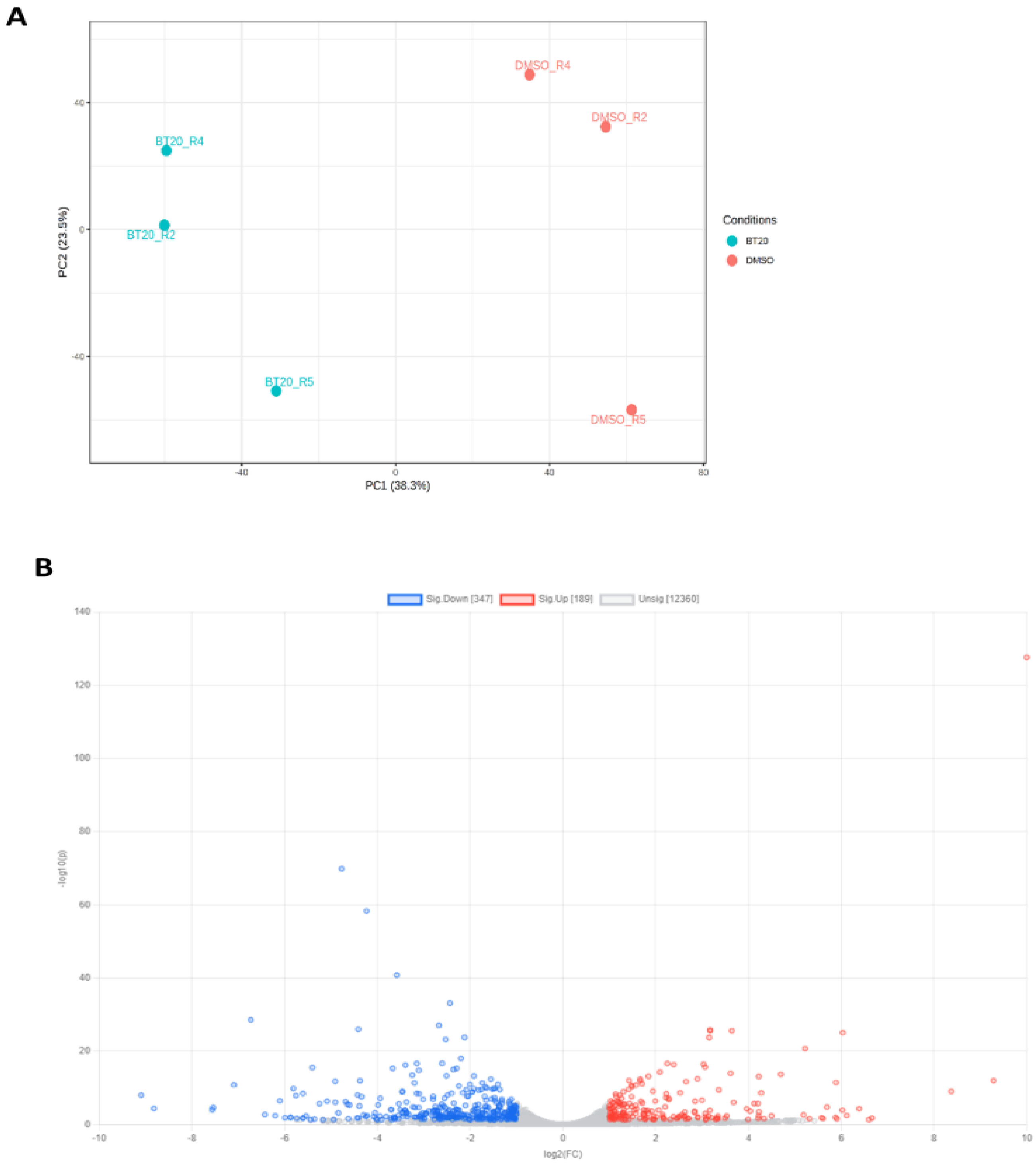


**Supplementary Figure 2. The control compound does not alter the transcriptomic profile of ductal organoids.** Shown are PCA (**A**) and volcano plots (**B**) obtained for ductal organoids treated with 20 µM BTYNB (BT20) or the vehicle DMSO**.** The patterns observed were similar to those observed for BT20 and CC40-treated organoids (shown in **Figure 4A-B**).

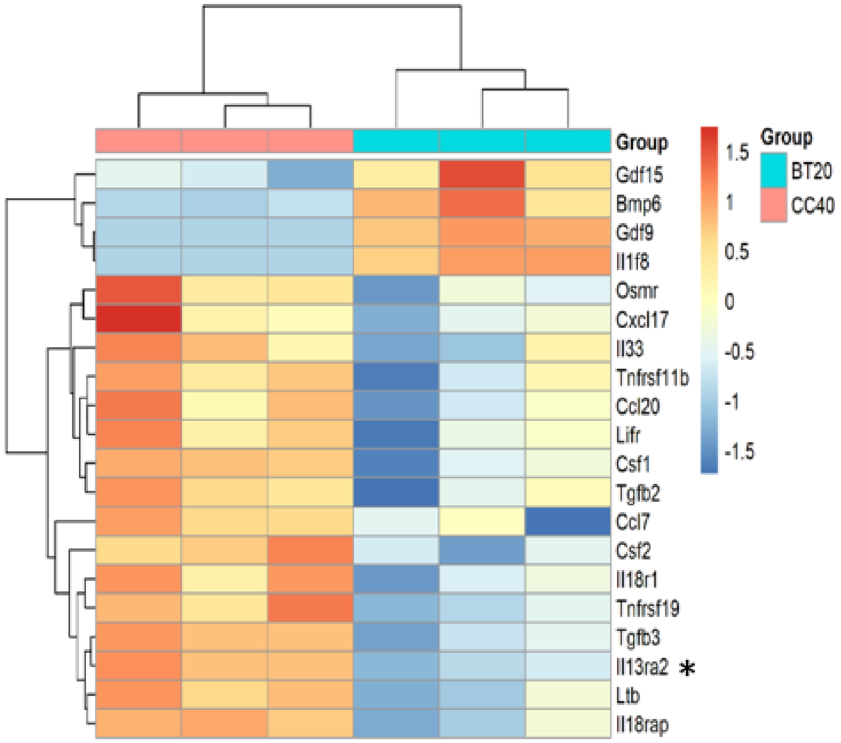


**Supplementary Figure 3. A significant downregulation of *Il13ra2* transcript is evident in BTYNB treated organoids**. Shown is a heatmap of gene enrichment in the cytokine-cytokine receptor interactions pathway. The *Il13ra2* transcript was the most downregulated among the genes enriched in this pathway (log fold change = -4.7098, FDR < 0.00001).

### SUPPLEMENTARY TABLES

| Case number | Age | Sex | Tumour grade | Tumour stage |
|---|---|---|---|---|
| 1 | 67 | M | G2 | pT3N1 |
| 2 | 80 | M | G2 | pT3N1 |
| 3 | 86 | M | G2 | pT2N0 |
| 4 | 66 | F | G2 | pT2N0 |
| 5 | 83 | M | G2 | pT2N1 |
| 6 | 67 | F | G1 | pT1cN0 |

**Supplementary Table 1: Patient characteristics and tumor grade.** Listed are the clinical data for six patients whose PDAC specimens were used for analysis of IMP1 expressions (Data shown in **Figure 1**).

| Sex of KPC+/- mice | Age (wks) at necropsy | Ascites | Pancreatic Tumor size and weight |
|---|---|---|---|
| male | 27 | +++ | 2.5 X1.5 X 0.7 cm<br>1.54 gm |
| male | 20 | - | 0.9 X 0.5 cm<br>0.529 gm |
| female | 23 | + | 2.0 X 1.5 cm<br>2.27 gm |
| female | 24 | + | 2 X1.3 X 0.5 cm<br>0.56 gm |
| female | 26 | +++ | 3.0 X 3.0 X 1.0 cm<br>3.54 gm |
| female | 31.5 | +++ | 1.7 X 1.1 cm<br>0.839 gm |

**Supplementary Table 2. Necropsy findings for KPC mice.** Shown is necropsy information for the KPC+/- mice used as source of pancreatic tissue in this study (**Figure 2**).

| Target gene | Forward sequence | Reverse sequence |
|---|---|---|
| Gkn3 | GCCTGTGTTCTGGCAAAGATGG | GGCTGGGTAAAACTGTGTAGGTC |
| Il13ra2 | AATGTTGGGAAGAGCCTCCA | GGCTGGCTCTATGTCAAGAAA |
| Vcam1 | GCTATGAGGATGGAAGACTC | ACTTGTGCAGCCACCTGAGA |
| IMP1 | GCCTCCATCAAGATTGCACC | ATTCTTCCCTGGGCCTTGAA |

**Supplementary Table 3. Primer sequences.** Listed are the primer sequences used for qPCR analysis.

| Biological process | Total | Expected | Hits | P.Value | FDR |
|---|---|---|---|---|---|
| Response to xenobiotic stimulus | 32 | 1.46 | 9 | 8.65E-06 | 0.00273 |
| Regulation of cell migration | 337 | 15.4 | 34 | 1.05E-05 | 0.00273 |
| Xenobiotic metabolic process | 26 | 1.19 | 8 | 1.33E-05 | 0.00273 |
| Sensory organ development | 326 | 14.9 | 33 | 1.33E-05 | 0.00273 |
| Cell migration | 680 | 31 | 55 | 1.87E-05 | 0.00286 |
| Negative regulation of multicellular organismal process | 275 | 12.5 | 29 | 2.09E-05 | 0.00286 |
| Positive regulation of MAPK cascade | 262 | 12 | 27 | 6.03E-05 | 0.00707 |
| Epithelial cell differentiation | 238 | 10.9 | 25 | 8.29E-05 | 0.00757 |
| Positive regulation of cell migration | 196 | 8.94 | 22 | 8.31E-05 | 0.00757 |
| Tube development | 378 | 17.2 | 34 | 0.000111 | 0.00913 |
| Endothelial cell migration | 86 | 3.92 | 13 | 0.000128 | 0.00958 |
| Morphogenesis of an epithelium | 353 | 16.1 | 32 | 0.000154 | 0.01 |
| Regulation of cell growth | 219 | 9.99 | 23 | 0.000159 | 0.01 |
| Tube morphogenesis | 270 | 12.3 | 26 | 0.000249 | 0.0146 |
| Organ morphogenesis | 665 | 30.3 | 50 | 0.000286 | 0.0156 |
| Tissue morphogenesis | 452 | 20.6 | 37 | 0.000375 | 0.0172 |
| Positive regulation of cell proliferation | 587 | 26.8 | 45 | 0.000383 | 0.0172 |
| Positive regulation of multicellular organismal process | 487 | 22.2 | 39 | 0.000405 | 0.0172 |
| Positive regulation of angiogenesis | 73 | 3.33 | 11 | 0.000432 | 0.0172 |
| Organ development | 2240 | 102 | 132 | 0.000457 | 0.0172 |

**Supplementary Table 4. BTYNB-mediated inhibition of IMP1 alters multiple biological processes**. Listed are the top 20 enriched biological processes (GO) that were affected by IMP1 inhibition ranked by the p value. The observed processes are associated mainly with regulation of cellular processes and organogenesis.

| Pathway | Total | Expected | Hits | P.Value | FDR |
|---|---|---|---|---|---|
| Metabolism of xenobiotics by cytochrome P450 | 41 | 2.12 | 10 | 3.13E-05 | 0.00632 |
| Complement and coagulation cascades | 43 | 2.23 | 10 | 4.88E-05 | 0.00632 |
| Chemical carcinogenesis | 44 | 2.28 | 10 | 6.04E-05 | 0.00632 |
| Cytokine-cytokine receptor interaction | 155 | 8.02 | 20 | 0.00012 | 0.00953 |
| Amoebiasis | 69 | 3.57 | 12 | 0.00018 | 0.0113 |
| Staphylococcus aureus infection | 28 | 1.45 | 7 | 0.00043 | 0.0193 |
| Rheumatoid arthritis | 55 | 2.85 | 10 | 0.00043 | 0.0193 |
| Ovarian steroidogenesis | 30 | 1.55 | 7 | 0.00067 | 0.0263 |
| MAPK signaling pathway | 215 | 11.1 | 22 | 0.00152 | 0.0529 |
| Mineral absorption | 35 | 1.81 | 7 | 0.00177 | 0.0556 |
| Thyroid hormone synthesis | 56 | 2.9 | 9 | 0.00207 | 0.059 |
| Drug metabolism - cytochrome P450 | 37 | 1.92 | 7 | 0.00248 | 0.0649 |
| IL-17 signaling pathway | 71 | 3.68 | 10 | 0.00327 | 0.0739 |
| TNF signaling pathway | 95 | 4.92 | 12 | 0.00336 | 0.0739 |
| PPAR signaling pathway | 50 | 2.59 | 8 | 0.00373 | 0.0739 |
| Ferroptosis | 30 | 1.55 | 6 | 0.00377 | 0.0739 |
| Steroid hormone biosynthesis | 31 | 1.6 | 6 | 0.00447 | 0.0826 |
| Glutathione metabolism | 43 | 2.23 | 7 | 0.00596 | 0.104 |
| PI3K-Akt signaling pathway | 249 | 12.9 | 22 | 0.00907 | 0.15 |
| Linoleic acid metabolism | 17 | 0.88 | 4 | 0.00978 | 0.154 |

**Supplementary Table 5. BTYNB-mediated inhibition of IMP1 alters multiple biological pathways.** Shown are the top 20 enriched biological pathways (KEGG) that were affected by IMP1 inhibition ranked by the p value. The pathways identified are related to pathological processes such as chemical carcinogenesis and inflammation, cell metabolism and cellular signaling.

### SUPPLEMENTARY DATA

**Supplementary Data 1. Differentially expressed genes in BTYNB treated cells.** Shown is the complete list of DEGs in BTYNB as compared to control treated organoids. The list of 535 transcripts with FDR < 0.05 and log fold change greater than 1 or less than -1 is also shown. Of these, 328 were downregulated in the BTYNB-treated organoids.

**Supplementary Data 2. Downregulated DEGs in BTYNB-treated organoids were also identified in the Pancreatic Cancer Database.** Each downregulated DEG (FDR < 0.05, log fold change < -1.5) was individually searched in the Pancreatic Cancer Database (http://pancreaticcancerdatabase.org/). The genes identified in the database are categorized based on expression in PDAC or precursor lesions and the changes observed in the expression level. *Il13ra2*, *Mmp9* and *Vcam1* (**in bold**) are all associated with invasive ductal adenocarcinoma.

### SUPPLEMENTARY METHODS

#### Mice

LSL-*Kras*$^{G12D}$ mice were kindly provided by Antonis Koromilas (Lady Davis Institute, Montreal, QC) and LSL-tdTomato mice were obtained from Jackson Laboratory (Strains: 007909). CluCreERT mice were described in detail elsewhere [1]. Sections from formalin fixed and paraffin embedded (FFPE) pancreatic tumors of LSL-KrasG12D/+;LSL-Trp53R172H/+;Pdx-1-Cre (KPC) mice crossbred at the UCSD animal care facility under animal care protocol S07095 were provided by the Lowy laboratory (See **Supplementary Table 2**). Sections from male and female B6/B6.129S4 mice were analyzed by immunohistochemistry as described. Acute pancreatitis was induced in adult male C57BL/6J mice (Jackson Laboratory) by intraperitoneal injection of caerulein (50 µg/kg; Sigma, cat. C9026) (or PBS as control) at hourly intervals eight times daily, for two consecutive days [2]. Mice were sacrificed the following day and the pancreas isolated for analysis.

#### Immunofluorescence staining of whole mount organoids

Pancreatic ductal organoids resuspended in Matrigel (Corning) were seeded on 15 mm circular glass coverslips (Chemglass inc.) and placed inside the wells of a 24-well culture plate (30 µl/well). Following 48-hour treatment with BTYNB, 1 µl of 10 mM Edu per 500 µl were added to each well and incubated at 37$^0$C for 1 hour. Organoids were then fixed with 10% buffered formalin on ice for 30 minutes and washed twice with PBS-0.1% Tween (PBST). Organoids were then permeabilized with 0.5% Triton X 100 in PBST for 20 minutes at RT, followed by washing twice with PBST and blocking with 1% bovine serum albumin (BSA) in PBST for 30 minutes. 300 µl of Edu staining solution (For 1 ml: 691 µl water, 100 µl 1M Tris pH 8.5, 1 µl 10 mM AlexaFluor488 Azide (Molecular probes #A10266), 8 µl 500 mM $CuSO_4$, 200 µl 500 mM L-Ascorbic acid (Sigma-Aldrich #A5960)) were added to each well and incubated for 1 hour at RT in the dark. After washing three times with PBST, organoids were incubated with antibody to cleaved caspase 3 (1:300, Cell Signaling Technologies, cat. 9661) in 1% BSA-PBST overnight at $4^0$C on a slow rocking platform in the dark, washed with PBST, and incubated with a corresponding Alexa Fluor secondary antibody for fluorescent-based detection and with DAPI for nuclear counter staining in 1% BSA-PBST for 2 hours at RT. Organoids were then cleared with fructose-glycerol clearing solution (60% (vol/vol) glycerol and 2.5M fructose in water) for 20 minutes and coverslips were inverted onto a microscope slide for imaging using the Zeiss LSM880 laser scanning confocal microscope.

**Synthesis of BTYNB**

Briefly, 1.05 g (5.51 mmol) of 5-bromothiophene-2-carbaldehyde were added to 0.5 g (3.67 mmol) of 2-aminobenzamide and heated to reflux overnight in 10 ml of ethanol. The crude mixture was evaporated, dissolved in dichloromethane and prepared as a solid deposit on silica gel. The deposit was loaded onto a BIotage cartridge and subjected to flash chromatography using the Biotage Isolera Prime under isocratic elution with 95%/5% dichloromethane/methanol. The fractions were pooled, evaporated and dried overnight to yield a crystalline white powder (1.55g, 88%). The control compound was synthesized using the same protocol with the 4-aminobenzamide and was obtained pure with a yield of 82%.

**Primary acinar cell isolation and three-dimensional culture**

Briefly, the pancreas was dissected, washed twice with ice-cold HBSS, minced into 1–5 mm fragments and digested with 10mg/ml collagenase P (25 minutes at 37 °C using a shaker). The digestion was terminated by adding an equal volume of ice-cold HBSS with 5% FBS. The digested pancreatic tissue was washed twice in 5 ml ice cold HBSS with 5% FBS, resuspended in the same medium and filtered using a 100 µM mesh filter. The filtrate containing the acinar cells was then carefully pipetted on top of 20 ml of an ice cold 30% FBS in HBSS solution and centrifuged at 1000 rpm for 2 minutes. The cells were gently suspended in Matrigel (Corning, cat. 356237) and plated at ~75% confluency in a 24-well plate (30 µl/well). After solidification at 37°C, culture media (RPMI 1640 complete medium with 1% FBS, 0.1 mg/ml trypsin inhibitor and 1 µg/ml dexamethasone), with or without TGFα (50 ng/ml; Preprotech, cat. 100-16A) were added to the wells [3, 4]. Acinar cells that transformed into the ductal phenotype were quantified by QuPath, and ductal cell size quantified by ImageJ or using OrganoID - a deep learning platform based on convolutional neural network [5] with cross verification that data generated by both quantification methods were comparable. Data processing was with GraphPad Prism (version 10.0.03). Statistical analyses were performed using 1-way ANOVA and statistical significance is indicated where applicable.

**Pancreatic ductal organoid culture**

Following collagenase dissociation (Collagenase type XI 0.012% (w/v) (Sigma), dispase 0.012% (w/v) (Gibco), FBS (Gibco) 1% in DMEM media (Gibco)) at 37°C, ducts were hand-picked under a dissection microscope and seeded in Matrigel (Corning) in a 24-well culture plate (30 µl/well). After solidification, 500 µl culture medium was added (Advanced DMEM (Invitrogen) supplemented with B27 (Invitrogen), 1.25 mM *N*-Acetylcysteine

(Sigma), 10 nM gastrin (Sigma) and growth supplements: 50 ng/ml EGF (Peprotech), 10% RSPO1-conditioned media (prepared in-house), 10% Noggin-conditioned media (prepared in-house), 100 ng/ml FGF10 (Peprotech) and 10 mM Nicotinamide (Sigma).

**CRISPR/Cas9 gene editing**

***Plasmid preparation*.** The LentiCRISPRv2-Puro plasmid was digested with the BsmBI (NEB) restriction enzyme and gel purified using the Biorad "Freeze and squeeze" DNA gel extraction kit, as per the manufacturer's instructions. The guide oligos (Forward 5'-caccg**AGAGGCTTACGAGAACGACG** -3' and Reverse 5'-aaac**CGTCGTTCTCGTAAGCCTCT**c -3') were prepared for plasmid insertion by phosphorylating the 5' ends with the T4 polynucleotide kinase enzyme (NEB), followed by annealing, initially at 95$^0$C, followed by gradual reduction of the temperature to 25$^0$C. The annealed oligos were inserted into the LentiCRISPRv2-Puro plasmid using the T4 DNA ligase (NEB) and used to transform the Stbl3 bacteria. The sequence was confirmed using the DNA sequencing services of the McGill University and Génome Québec Innovation Centre.

**Generation and selection of IMP1-silenced LMP cells**

All cells were maintained as a frozen stock and generally cultured *in vitro* for up to 4 weeks only prior to use in the *in vivo* experiments, in order to minimize genetic drifts and changes to their metastatic phenotypes. They were expanded in a humidified incubator at 37°C with 5% $CO_2$ in DMEM medium (Wisent), supplemented with 4 mM L-glutamine, 4.5 g/L glucose, a solution of 100 U/ml penicillin and 100 µg/ml streptomycin (Sigma), 2 g/L sodium-pyruvate and 10% fetal bovine serum (FBS; Wisent). LMP cells maintained in DMEM/F12 medium supplemented with 10% FBS were cultured to 60-80% confluency and transfected with the LentiCRISPRv2-Puro plasmid (prepared as described above) containing the IMP1 guide or an irrelevant sequence (Rosa26, known to have no specific importance in mice) as control, using Lipofectamine2000 as per the manufacturer's (ThermoFisher scientific) instructions. The transfected cells were incubated at $37^0$C in full media overnight, at which time the medium was replaced with medium containing 2µg/ml puromycin for a further incubation of 48 hours. The surviving cells were dispersed using 0.025 % trypsin in $Ca^{++}$ and $Mg^{++}$ Free PBS-EDTA, and single cells plated in 96 well plates and incubated for up to 3 weeks in complete medium. Individual clones were selected, expanded and Western blotting performed using an antibody to IMP1 (Cell Signaling clone: D33A2, cat- 8482) to identify clonal populations in which IMP1 expression was silenced. The silenced clones and control cells were expanded, and silencing confirmed by DNA sequencing, using the services of the McGill University and Génome Québec Innovation Centre.

**Histology, immunohistochemistry and immunofluorescence**

***Histology.*** Formalin fixed tissue obtained from patients (**Supplementary Table 1**) was processed, embedded in paraffin, and cut into 5 µm sections. Hematoxylin and Eosin (H&E) (ThermoFisher Scientific, cat. 7221, 7111) staining was performed according to the clinical laboratory standard. Two areas of normal acini, ADM and PDAC were selected from each case for construction of a tissue microarray (TMA).

***Immunohistochemistry.*** Immunohistochemistry to detect IMP1 in PDAC tumors and precursor lesions was performed with the help of the Histopathology Platform of the RI-MUHC on formalin fixed paraffin embedded (FFPE) pancreatic tissue removed from Trp53+/-/LSL-KRAS*G12D* X Pdx1-Cre+/- (KPC) B6/B6.129S4 mice bred at the UCSD animal facility and maintained as per AUP S07095 and on the TMA prepared as described above. Five µm sections of the TMA were prepared from the paraffin blocks, deparaffinized in xylene and rehydrated in graded ethanol. Antigen retrieval was performed by heating the sections in boiling sodium citrate buffer (Sigma-Aldrich, cat. C-9999) for 20 minutes. After blocking with 3% hydrogen peroxide and bovine serum albumin (BSA), tissues were incubated with anti-IMP1 antibody (Cell Signaling Technologies, cat. 2852, diluted 1:1000) at 4°C overnight, washed and incubated with horseradish peroxidase (HRP)-conjugated secondary antibodies. The color was developed using diaminobenzidine (DAB) substrate (Sigma-Aldrich, cat. D-7304) and slides were counterstained with hematoxylin.

IHC on FFPE sections of KPC–derived PDAC (4 µm-thick) was performed using the Discovery Ultra instrument (Roche). Sections were deparaffinized, rehydrated and incubated for 40 min at $37^0$C with anti-IMP1 antibody (Thermo Fisher, PA5-44886, 1:50) following antigen retrieval treatment (56 min in EDTA buffer). This was followed by a 20 min incubation at room temperature with OmniMap anti-rabbit-HRP (Roche 760-4311). Color was detected using the ChromoMap DAB kit (760-4304, Roche) and sections counterstained with hematoxylin, dehydrated, cleared and cover slipped. The slides were digitally scanned using an Aperio scanner (Leica, Aperio Turbo 2T) for morphometric analysis with the Imagescope software.

***Immunofluorescence***. Immunofluorescence staining was performed using primary antibodies against IMP1 (1:500, Cell Signaling Technologies, cat. 2852), cytokeratin-19 (CK19; 1:500, DSHB, TROMAII), and alpha-smooth muscle actin (SMA, 1:2000, Sigma-Aldrich cat. A2547). Corresponding Alexa Fluor dyes were used for fluorescent detection and DAPI was used for nuclear counter staining. Images were captured on the Zeiss LSM880 laser scanning confocal microscope.

***Cell proliferation assays***. Tumor cell proliferation was measured using the colorimetric MTT assay. LMP cells in DMEM containing 10% FBS were seeded in each well of a 96-well plate and incubated overnight. Cells were then treated with DMSO (vehicle), BYTNB, or a control inactive compound (CC) at the indicated concentrations. The media was refreshed every two days. At each time point, cells were incubated for 3 hours with 10µl of 5 mg/ml MTT Reagent at 37°C. After incubation, crystalline formazan was solubilized with 100µl of DMSO for 15 min at RT. Absorbance was measured at OD=570 using a microplate reader. Where indicated, cell proliferation was also measured over a period of 3 days using the Trypan blue viability dye and viable cells enumerated manually using a microscope.

**Immunoblotting**

Immunoblotting was performed essentially as we described in detail previously[6]. Briefly, cultured cells were washed with TBST (ThermoFisher Scientific, USA) and lysed in a lysis buffer containing 50mM Tris (pH 7.4), 1mM EDTA, 150mM NaCl, 1% (w/v) NP-40, 2mM Na3VO4, 5mM NaF, 0.25% sodium deoxycholate and a protease inhibitor cocktail (Roche cOmplete Mini-Mississauga, ON, CA, Sigma-Aldrich- Oakville, ON. Canada) then scraped and centrifuged at 12000 x g for 5 minutes. Proteins in the supernatant were quantified, separated on 10% SDS-PAGE gels and transferred onto nitrocellulose membranes. The membranes were blocked with 3% BSA in DPBS containing 0.1% Tween 20 for 30 minutes, followed by incubation overnight at $4^0$C with the primary antibody in DPBS+ 0.1% Tween 20 and then for 1 hr at RT with HRP-conjugated anti-rabbit or anti-mouse immunoglobulin secondary antibodies (Jackson ImmunoResearch-West Grove, PA, USA), as appropriate. Signal detection and densitometry were performed using ImageQuant Las4000.

**Experimental liver metastasis**

Experimental liver metastases were generated by intrasplenic/portal injections of $10^5$ LMP cells followed by splenectomy as we previously described [7]. Animals were euthanized 21 days later, and visible metastases on the surface of the liver were enumerated and sized without prior fixation.

**RNA sequencing**

We treated KrasG12D-expressing ductal organoids with 20 µM BTYNB (BT20), vehicle (DMSO) or the inactive control compound at 20 or 40 µM (CC40, for increased specificity). Total RNA from these cells was extracted using the PureLink™ RNA Mini Kit (Invitrogen, cat. 12183020) as per the manufacturer's instructions and sent to the McGill University and Génome Québec Innovation Centre for paired-end read, next generation sequencing of RNA (100 bp in sequence length) using the Illumina NovaSeq 6000 platform. The fastq data were obtained from the facility and were analyzed using the RNAseq pipeline described in detailed elsewhere [8, 9].

Briefly, quality analysis of the raw data was checked with FASTQC (version 11.3) and adapter related sequences were removed using Trim Galore (version 0.6.6) (https://www.bioinformatics.babraham.ac.uk/projects/). The genome sequence of mouse (*Mus musculus*) and its annotation file (Mus_musculus.GRCm39.107) were obtained from ENSEMBL (https://www.ensembl.org/). Reads were aligned to the mouse genome with HISAT2 (version 2.2.0) [10] and read counts were obtained using HTSeq (version

2.0.2) [11], where the intersection-strict mode was applied. Exploratory analysis was performed with ExpressAnalyst. Low variance (15%) and low abundance ($n < 4$) counts were filtered. Data were normalized as trimmed mean of M-values and sample distribution was visualized by principal component analysis. Differential gene expression analysis was performed using edgeR [12] and differentially expressed genes (DEGs) with FDR values of less than 0.05 were accepted as statistically significant. Overrepresentation enrichment network and heatmap clustering were performed on the data. Biological processes and pathways were considered significant when the p value determined by the enrichment analysis was less than 0.05. Heatmaps were generated with the pheatmap function in R (version 4.2.2). Conversion of NCBI gene IDs to Ensemble gene IDs was performed with the DAVID Gene ID Conversion Tool [13].

**Quantitative PCR (qPCR)**

Expressions of genes of interest, identified by the RNAseq analysis, were validated using qPCR. The cDNA was synthesized from extracted RNA using M-MLV Reverse Transcriptase (Invitrogen) and the PCR performed with FastStart Universal SYBR Green Master (Roche) and the primer sequences listed in **Supplementary Table 3** using the 7500 Real-Time PCR System (Applied Biosystems™). Expression levels were compared using the 2(-Delta Delta C(T)) method [14]. Expression levels in organoids derived from $Kras^{G12D}$ mice were compared to wild type organoids using the same method and three replicates for each organoid type.
